# PCB126 exposure revealed alterations in m6A RNA modifications in transcripts associated with AHR activation

**DOI:** 10.1101/2020.07.02.182865

**Authors:** Neelakanteswar Aluru, Sibel I Karchner

**Affiliations:** Biology Department and Woods Hole Center for Oceans and Human Health Woods Hole Oceanographic Institution, Woods Hole, MA 02543

**Author notes:** Corresponding author: Neelakanteswar Aluru, Biology Department, MS-32, Woods Hole Oceanographic Institution, Woods Hole, MA 02543 U.S.A.

**Keywords:** *dioxin-like PCBs*, development, zebrafish, epitranscriptomics, m6A, MeRIP

## Abstract

Chemical modifications of proteins, DNA and RNA moieties play critical roles in regulating gene expression. Emerging evidence suggests these RNA modifications (epitranscriptomics) have substantive roles in basic biological processes. One of the most common modifications in mRNA and noncoding RNAs is *N*^6^-methyladenosine (m6A). In a subset of mRNAs, m^6^A sites are preferentially enriched near stop codons, in 3’ UTRs, and within exons, suggesting an important role in the regulation of mRNA processing and function including alternative splicing and gene expression. Very little is known about the effect of environmental chemical exposure on m6A modifications. As many of the commonly occurring environmental contaminants alter gene expression profiles and have detrimental effects on physiological processes, it is important to understand the effects of exposure on this important layer of gene regulation. Hence, the objective of this study was to characterize the acute effects of developmental exposure to PCB126, an environmentally relevant dioxin-like PCB, on m6A methylation patterns. We exposed zebrafish embryos to PCB126 for 6 hours starting from 72 hours post-fertilization and profiled m6A RNA using methylated RNA immunoprecipitation followed by sequencing (MeRIP-seq). Our analysis revealed 117 and 217 m6A peaks in the DMSO and PCB126 samples (FDR 5%), respectively. The majority of the peaks were preferentially located around the 3’UTR and stop codons. Statistical analysis revealed 15 m6A marked transcripts to be differentially methylated by PCB126 exposure. These include transcripts that are known to be activated by AHR agonists (e.g., *ahrra, tiparp, nfe2l2b) as well as others that are important for normal development (vgf, cebpd, foxi1). These results suggest that* environmental chemicals such as dioxin-like PCBs could affect developmental gene expression patterns by altering m6A levels. Further studies are necessary to understand the functional consequences of exposure-associated alterations in m6A levels.

## Introduction

Epigenetic regulation of gene expression has been shown to play critical roles in basic biological processes in vertebrates. Enormous strides have been made in our understanding about the role of DNA methylation and chromatin modifications in gene regulation. Recent studies have demonstrated another layer of epigenetic regulation at the RNA level, where internal chemical modifications of mRNA have been shown to affect their stability, processing, localization and splicing. There are over 160 different RNA modifications that have been identified and their roles in embryonic development, nervous system function and multigenerational effects have been demonstrated.

One of the most common modifications in mRNA and noncoding RNAs is *N*^6^-methyladenosine (m6A). In a subset of mammalian mRNAs, m^6^A sites are preferentially enriched near stop codons, in 3’ UTRs, and within long internal exons with a consensus sequence of RRACH (R=G or A; H=A, C or U), suggesting an important role in the regulation of gene expression. m6A has been shown to regulate RNA processing, including RNA splicing (Kasowitz et al. 2018; Tang et al. 2018), nuclear export (Roundtree et al. 2017), RNA degradation (Zheng et al. 2013), and translation (Huang et al. 2018; Shi et al. 2017). The enzymatic machinery involved in writing, reading and removing the m6A modifications have been identified. A multicomponent writer complex consisting of the catalytic subunit methyltransferase like 3 (METTL3) is responsible for m6A RNA methylation, while fat mass and obesity-associated protein (FTO) and α-ketoglutarate-dependent dioxygenase alkB homolog 5 (ALKBH5) have been identified as the m6A erasers (Meyer and Jaffrey 2017). Additionally, m6A can be recognized by multiple RNA-binding proteins, such as the YTH domain family proteins and the heterogeneous nuclear ribonucleoprotein (HNRNP) family (Meyer and Jaffrey 2017; Roundtree et al., 2017; Zhou et al. 2019). Gene deletion studies have revealed that loss of *mettl3* in mice is embryonic lethal and leads to complete loss of m6A in polyadenylated RNA (Geula et al. 2015). In contrast, *fto* knockout mice develop normally and show elevated m6A levels selectively in the 5’UTR (Jia et al. 2011) but recent results suggests that *fto* non-specifically targets m6A (Mauer and Jaffrey 2018; Zaccara et al. 2019). In addition, deletion of a YTH domain containing protein, *ythdf2* has been shown to increase the half-lives of mRNAs compared to those lacking m6A (Zhao et al. 2017). These studies demonstrate that m6A RNA methylation is under the regulation of these three important group of proteins. There is growing evidence that these proteins play an important role in a number of physiological processes including stress response (Batista et al. 2014; Chen et al. 2015; Engel et al. 2018; Fischer et al. 2009; Schwartz et al. 2014; Zhang et al. 2017; Zhao et al., 2017). However, there is very limited information about the impact of environmental chemicals on m6A methylation patterns and the associated functional consequences on gene expression patterns. Recent studies have demonstrated that environmental chemicals such as arsenite affect RNA methylase and demethylase protein expression (Bai et al. 2018; Cayir et al. 2019; Chen et al. 2019; Gu et al. 2018; Zhao et al. 2019). These studies have also measured global m6A RNA content using an ELISA-based method. However, this approach does not provide any information about the location of the m6A modifications.

Even though the existence of m6A RNA modification has been known for a long time (Desrosiers et al. 1974), recent advances in the transcriptome-wide m6A quantification methods (Dominissini et al. 2013; Linder et al. 2015; Meyer et al. 2012) have yielded unprecedented insights into the location of m6A in RNAs. To date, majority of the studies have relied on immunoprecipitation of methylated RNAs using m6A-recognizing antibodies such as methylated RNA immunoprecipitation followed by sequencing (MeRIP-seq) (Meyer et al., 2012). Using MeRIP-seq, we tested the hypothesis that developmental exposure to environmental chemicals causes alterations in m6A RNA methylation patterns.

The objective of this study was to characterize the acute effects of developmental exposure to PCB126, an environmentally relevant dioxin-like PCB, on m6A methylation patterns. The experiments reported here were conducted in zebrafish, a well-established and widely used model for developmental toxicology (Behl et al. 2019; Nishimura et al. 2016) and human diseases (Bradford et al. 2017; Grunwald and Eisen 2002. Zebrafish embryos were exposed to PCB126 for 6 hours starting from 72 hours post-fertilization and profiled m6A RNA using MeRIP-seq, as well as assessing changes in mRNA expression and splicing. As many of the commonly occurring environmental contaminants alter gene expression profiles and have detrimental effects on physiological processes, it is important to understand the effects of exposure on this important layer of gene regulation.

## Materials and methods

### Experimental animals

The wild-type AB strain of zebrafish was used in this study. All experiments were conducted as per protocols approved by the Animal Care and Use Committee of the Woods Hole Oceanographic Institution. Freshly fertilized eggs were obtained from breeding of multiple tanks with 20-25 male and female fish.

### Developmental exposure to PCB126

Zebrafish embryos were exposed to 10 nM PCB126 (3,3’,4,4’,5-pentachlorobiphenyl, 99.2% purity; UltraScientific, RI, USA) or DMSO (solvent control; 0.01%) starting at 72 hours post-fertilization (hpf) for 6 hours. This duration of exposure was chosen based on differential gene expression observed in previous studies. In addition, short term exposure allows us to characterize primary changes in the m6A patterns. Each treatment consisted of four biological replicates with 48 embryos per replicate. Embryos were maintained in glass petri dishes at 28 ± 0.5°C at a density of 1 embryo per mL in 0.3X Danieau’s solution (pH 7.2). At the end of the exposure, embryos were thoroughly rinsed with 0.3X Danieau’s solution and sampled for MeRIP sequencing.

### Total RNA isolation

DNase-free total RNA was isolated using the BioRad Aurum kit following manufacturer’s instructions. Quantity and quality of total RNA was determined using the Qubit RNA high sensitivity assay kit (Thermo Fisher Scientific) and Agilent Bioanalyzer, respectively. All samples had an RNA integrity number above 9.3.

### MeRIP sequencing (MeRIP-seq)

m6A RNA was immunoprecipitated with an anti-m6A antibody (rabbit polyclonal; Synaptic Systems, Germany; catalog #202 003; RRID: AB_2279214). This antibody has been extensively used for m6A RNA immunoprecipitation (Dominissini et al., 2013; Engel et al., 2018; Zeng et al. 2018; Zhong et al. 2018), including zebrafish (Zhao et al., 2017). MeRIP-seq was conducted following a recently published protocol using low quantities of total RNA (Zeng et al., 2018). Briefly, the protocol involves RNA fragmentation, m6A immunoprecipitation and sequencing library preparation. For each sample, 18 ug of total RNA was chemically fragmented in 20 uL volume with the RNA fragmentation reagent at 70°C (Thermo Fisher Scientific Inc.). The fragmentation reaction was stopped by adding a stop solution and precipitated overnight at −80°C with 3M sodium acetate, glycogen and 100% ethanol. The precipitated RNA was pelleted by centrifugation, washed with 75% ethanol and resuspended in molecular grade water. The size distribution of fragmented RNA was assessed using the Bioanalyzer.

m6A RNA was immunoprecipitated by incubating the fragmented RNA with 5ug of anti-m6A antibody-magnetic bead mixture. This mixture was prepared by incubating the antibody with 30 uL each of protein A and G magnetic beads in immunoprecipitation buffer for at least 6 hours at 4°C. The antibody-bead mixture was washed with IP buffer and incubated with fragmented RNA for 2 hours at 4°C. RNasin Plus RNase inhibitor was added to the incubation mix to prevent RNA degradation. Following incubation, the antibody-bead mixture was washed twice with IP buffer, followed by low and high salt buffers for 10 minutes each at 4°C. The m6A-enriched fragmented RNA was eluted from the beads and purified using the RNeasy mini kit (Qiagen). The quality of the m6A-enriched RNA was assessed with the Bioanalyzer. m6A and input RNA libraries were prepared using the SMARTer stranded total RNA-seq kit (Clontech) and sequenced on the Illumina NextSeq500 platform (75bp, paired end reads).

### MeRIP sequencing (MeRIP-seq) data analysis

#### Mapping

Data analysis (Fig. 1) was carried out following previously established protocols for analysis of genome-wide m6A patterns (Dominissini et al., 2013). Raw reads from individual samples were pre-processed using Trimmomatic to remove adapters and low quality reads. Ribosomal reads were removed from downstream processing by mapping the pre-processed reads to the ribosomal sequences. The zebrafish ribosomal sequences were downloaded from the Silva database (https://www.arb-silva.de/). Ribosomal-free reads were mapped to the zebrafish genome (GRCz11 version 96) using the STAR aligner version 2.6.0d (Dobin et al. 2013). The resulting unique reads were used for peak calling.

**Figure 1.**
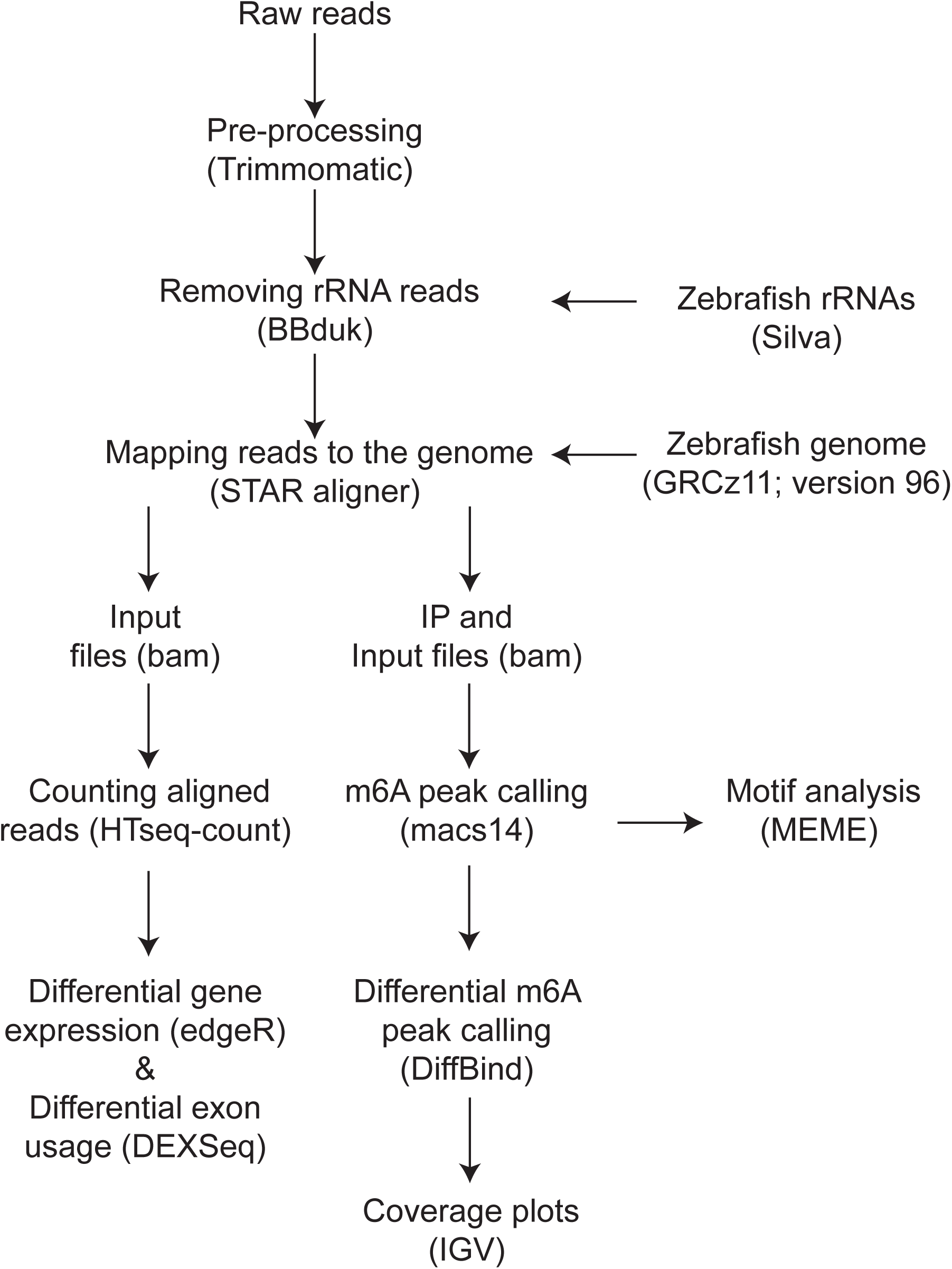
Overview of the MeRIP analysis pipeline. Raw reads were trimmed and ribosomal RNA reads were filtered prior to mapping to the genome and m6A peak calling, followed by statistical analysis. Integrated Genome Viewer (IGV) was used to visualize the m6A peaks. Input reads were used for differential gene expression and exon usage analysis. The detailed description of the analysis is provided in the methods section.

#### Peak calling and visualization

We used Model-based Analysis of ChIP-Seq (MACS) algorithm (version 1.4.2) for identification of the m6A peaks. The immunoprecipitated (m6A) and input (fragmented RNA) samples were used as treatment and control input files, respectively. The effective genome size (--gsize) was set to 117,608,789. The coverage data (wiggle files) was normalized to obtain comparable read density between input and IP samples and visualized using the UCSC genome browser.

#### Motif search and peak annotation

*De novo* motif finding was done using MEME (Bailey et al. 2009). The FASTA file of the 50 bp region flanking the peak summits was used as input and the top 3 consensus motifs were retrieved. CentriMO was used for visualization of the positional distribution of the best match of the m6A motif in the 300 bp centered around the peak summits.

#### Statistical analysis

Differential methylation analysis was done following the guidelines recommended for MeRIP analysis (McIntyre et al. 2020). We used DiffBind, a bioconductor package, for differential m6A peak calling (Stark and Brown 2011).

### Gene expression profiling

The input RNA samples were used for differential gene expression analysis. The number of reads mapped to the annotated regions of the genome were obtained with HTSeq-count (Anders et al. 2015). Statistical analysis was conducted using edgeR, a Bioconductor package (Robinson et al. 2010). We used the quasi-likelihood model in edgeR (glmQLFTest) to perform differential gene expression analysis. Only genes with a false discovery rate (FDR) of less than 5% were considered to be differentially expressed. Annotation of the differentially expressed genes was done using BioMart (Smedley et al. 2015).

### Differential exon usage analysis

To obtain the differentially used exons (DUEs) we used DEXSeq (Anders et al. 2012) and compared the exon usage between DMSO and PCB126 exposed groups in input RNA samples. DUEs are defined as a change in exon usage where exon usage is defined as the ratio of reads mapping to a particular exon vs reads mapping anywhere else in a gene (Anders et al., 2012). RNAseq data from the input samples were used for estimating DUEs. We considered exons to be differentially used when their adjusted p-value was below 0.05 (Benjamini Hochberg correction).

## Results

All the raw and processed data from this study have been deposited into Gene Expression Omnibus (Accession number GSE153436).

### m6A methylation profiles in zebrafish

MeRIP-sequencing of zebrafish embryos revealed highly conserved characteristics of m6A RNA methylation. For instance, m6A RNA methylation is predominantly found in the 3’UTR. The metagene plot of a subset of the m6A peaks shows the characteristic peak in RNA methylation around the end of the coding sequence and beginning of the 3’UTR (Fig. 2A). The pie chart representation of all the m6A peaks show that 51% of the peaks are found in the 3’UTR (Fig. 2B). Motif search of the m6A peaks identified previously established RRACH consensus sequence observed in vertebrates (E-value = 5.2E-35; Fig. 2C).

**Figure 2.**
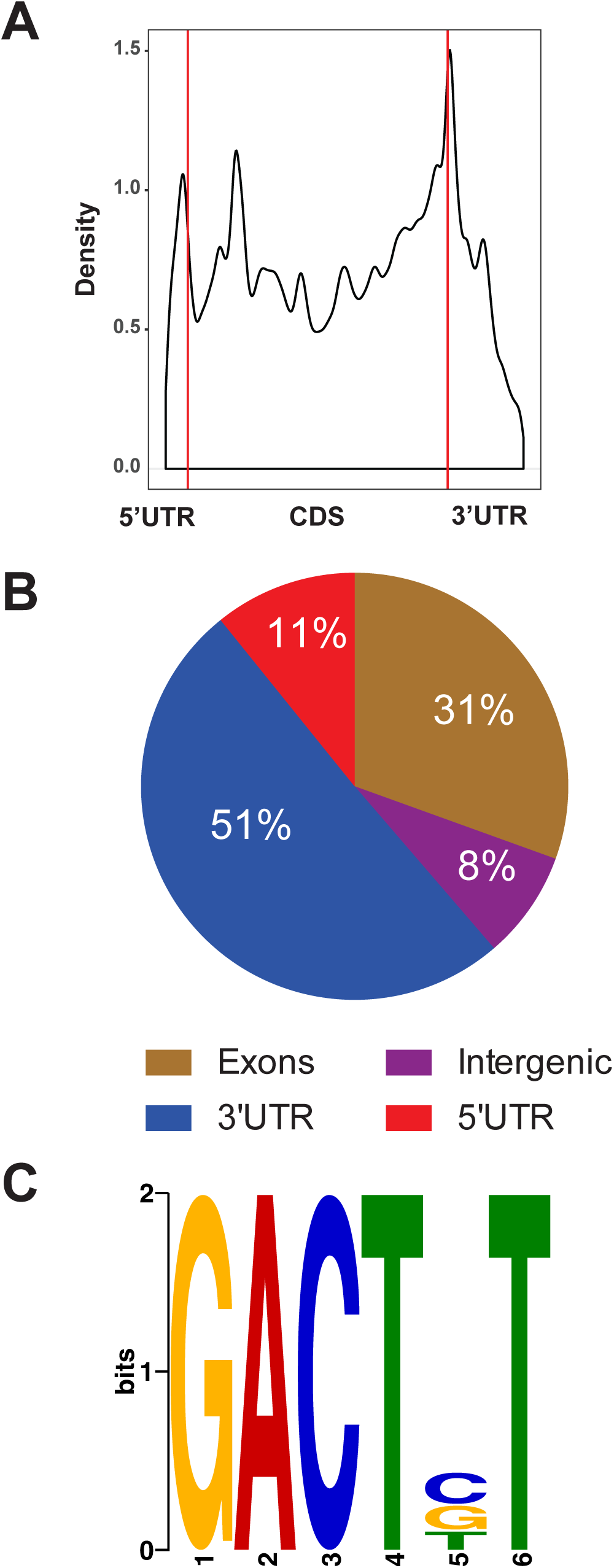
m6A RNA methylation landscape in zebrafish embryos. (A) Metagene profiles of m^6^A distribution across the transcriptome in control samples. (B) Distribution of m6A peaks in the different transcript regions (5’ and 3’UTR, exons and introns). (C) Consensus sequence motif for m^6^A RNA methylation identified in control samples.

The number of m6A peaks among the DMSO replicates range from 156 to 875 (FDR 5%). Among them 117 peaks were common to all biological replicates (Fig. 3A). Gene ontology (GO) term analysis of the common peaks revealed enrichment of biological process terms – nervous system development (GO:0007399), regulation of alternative mRNA splicing via spliceosome (GO:0000381) and RNA-dependent DNA biosynthetic process (GO:0006278) and molecular function terms – methyl CpG binding (GO:0008327) and first spliceosomal transesterification activity (GO:0000384) (Fig. 3B).

**Figure 3.**
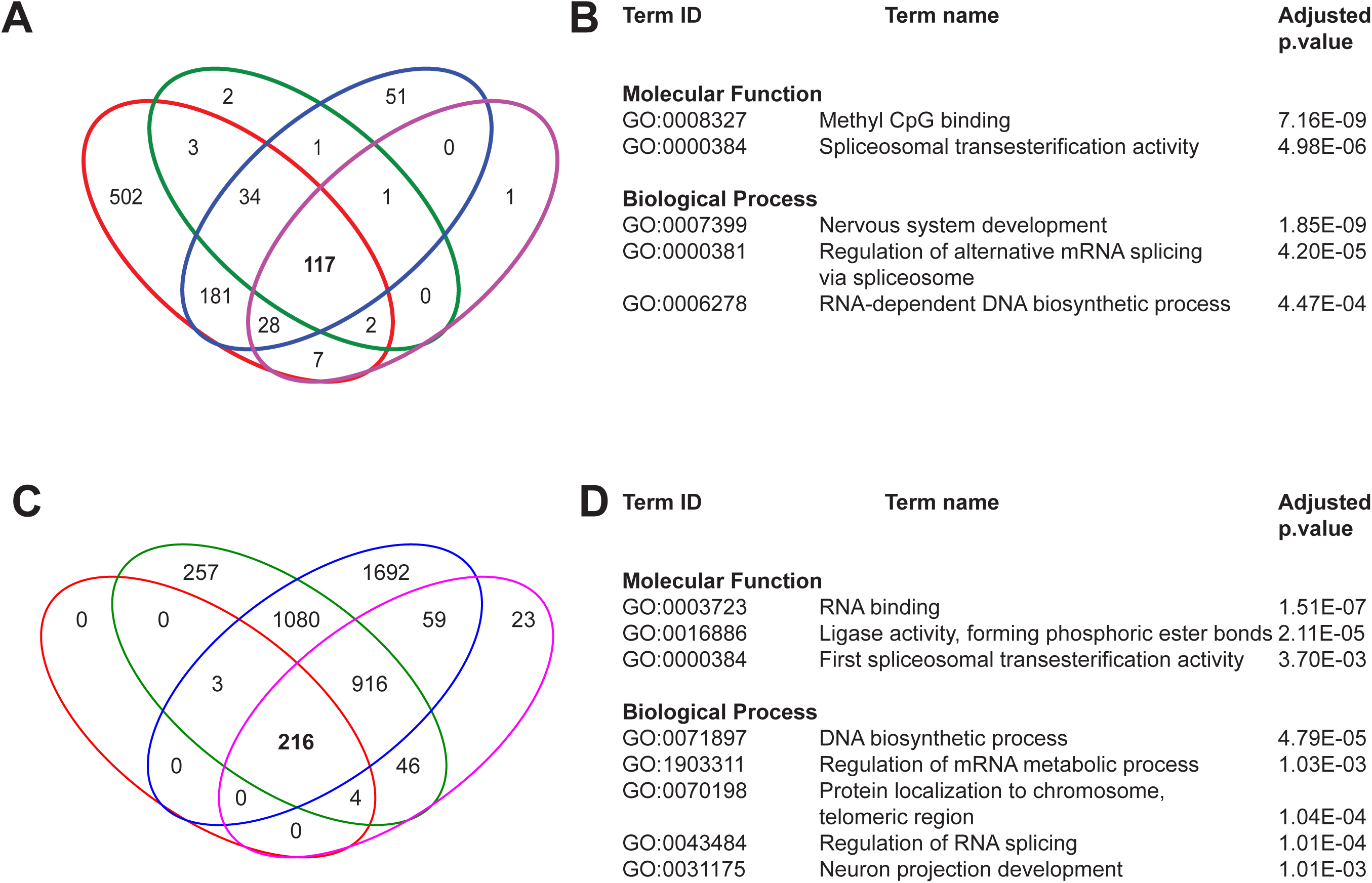
m6A RNA modification patterns in DMSO and PCB126 samples. (A and C) Venn diagram showing the unique and shared m6A peaks among biological replicates in DMSO and PCB126 groups. (B and D) Gene Ontology analysis (Biological Process and Molecular Function) of m6A peaks common to all 4 biological replicates in DMSO and PCB126.

Similarly, the number of m6A peaks among the PCB replicates range from 224 to 3971 (FDR 5%). Among them 217 peaks were common to all biological replicates (Fig. 3C). Gene ontology (GO) term analysis of the common peaks revealed enrichment of biological process terms – DNA biosynthetic process (GO:0071897), regulation of mRNA metabolic process (GO:1903311), protein localization to chromosome, telomeric region (GO:0070198), regulation of RNA splicing (GO: 0043484) and neuron projection development (GO:0031175) and molecular function terms – RNA binding (GO:0003727), ligase activity (GO:0016886) and first spliceosomal transesterification activity (GO:0000384) (Fig. 3D).

### PCB126 exposure induced changes in m6A RNA methylation

Principal component analysis (PCA) revealed one of the treatment biological replicates as an outlier (Supplemental figure 1). We omitted this replicate from statistical analysis. Based on the comparison between peaks that were present in both PCB126 (3 replicates) and DMSO (4 biological replicates) samples, we observed 15 differentially methylated peaks (DESeq2, 5%FDR), and 11 of these showed higher methylation in response to PCB126 (Fig. 4A). The genomic coordinates, read counts and the statistical significance of the differentially methylated peaks is provided in supplemental information (Differential_m6A_methylation_DiffBind.xlsx). These peaks include aryl hydrocarbon receptor repressor a (*ahrra*), TCDD inducible poly(ADP-ribose) polymerase (*tiparp*), pleckstrin homology, MyTH4 and FERM domain containing H3 (*plekhh3*), nuclear factor, erythroid 2-like 2b (*nfe2l2b*), protein phosphatase 1 regulatory subunit 18 (*ppp1r18*), retinitis pigmentosa GTPase regulator a (*rpgra*), znf648, pleckstrin homology domain-containing family G member 4B (*plekhg4b*, si:dkey-65j6.2), FP236157.6 (lincRNA), collagen type XXIV alpha 1 chain (CABZ01071903.1) and one unannotated region. Four differentially methylated peaks were reduced in response to PCB126 exposure and these include VGF nerve growth factor inducible (*vgf*), CCAAT enhancer binding protein delta (*cebpd*), si:dkey-175g6.2 (orthologous to human VGF) and forkhead box i1 (*foxi1*). Representative m6A peak coverage plots from *ahrra* (Fig. 4B) and *vgf* (Fig. 4C) are shown as examples.

**Figure 4.**
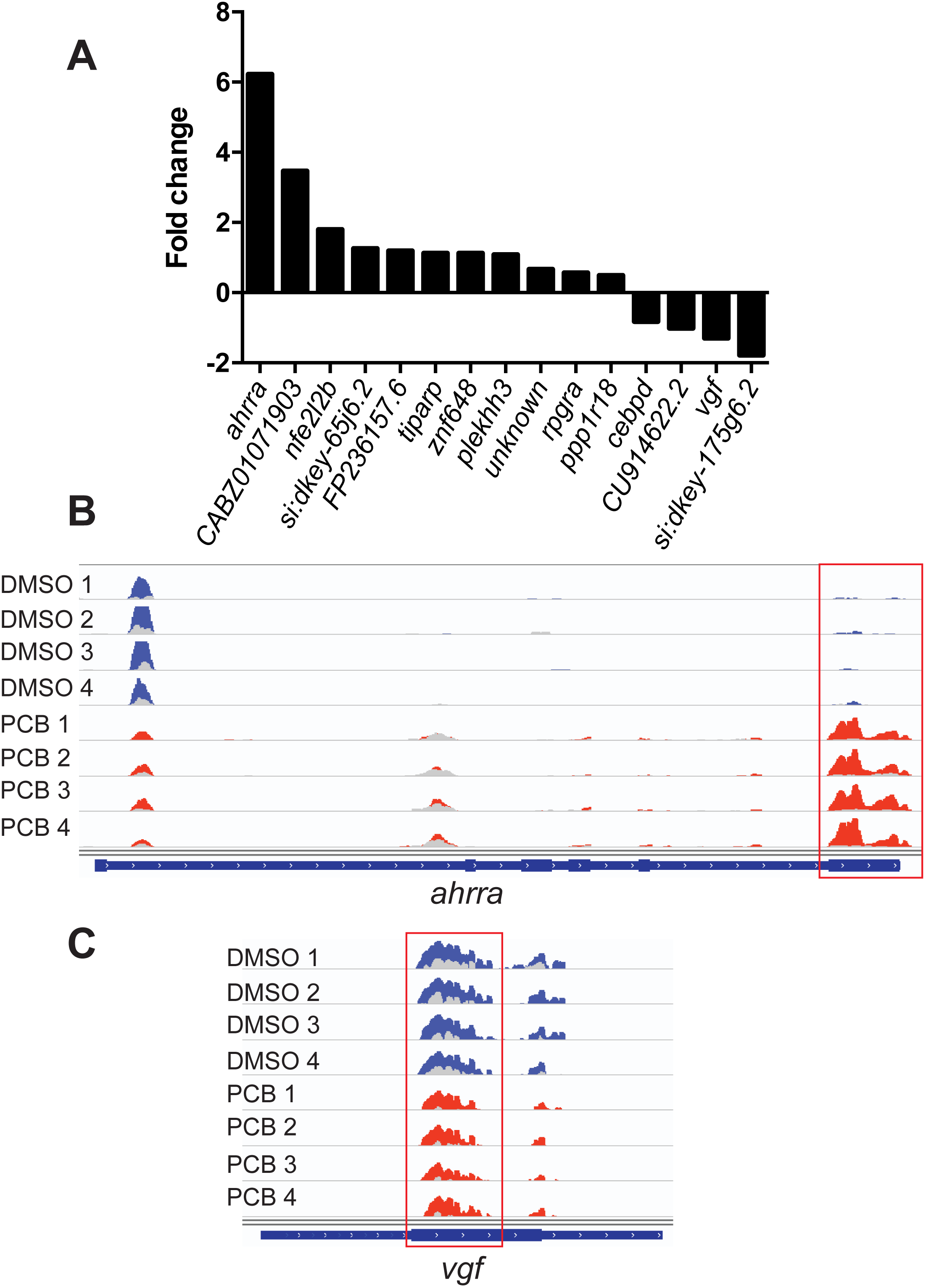
PCB126-induced alterations in m6A methylation patterns. (A) Histogram showing the transcripts with differentially methylated m6A peaks. (B) Representative m6A peak coverage plots in DMSO and PCB126 samples. Each track is an individual biological replicate (n=4). Differentially methylated m6A peak is highlighted with a red square. The input sample tracks (light color) are overlaid over the immunoprecipitated (IP) sample. DMSO and PCB126 tracks are shown in blue and red color, respectively.

### PCB126 exposure induced differential gene expression and exon usage

RNAseq analysis of the input RNA revealed differential expression of 272 genes in response to PCB126 exposure, with 85 and 187 up and downregulated genes, respectively (Fig. 5). The complete list of differentially expressed genes is provided in Supplementary Material (MeRIP-InputRNA-DGE.xlsx). The upregulated genes are enriched in the GO biological process term response to xenobiotic stimulus, and GO molecular function terms such as cofactor binding, iron ion binding, intermediate filament binding and oxidoreductase activity (Fig. 5B). The upregulated genes include prototypical AHR target genes (*cyp1a, cyp1c1, cyp1c2, cyp1b1, ahrra, foxf2a)*. The downregulated genes are enriched in GO terms such as response to peptide (biological process), organic ion transmembrane transporter activity, oxidative phosphorylation uncoupler activity and steroid hormone receptor binding (Fig. 5C). Differential exon usage analysis revealed two transcripts (protocadherin10 and solute carrier family 66 member 1) with significant differences in exon usage. Both of these transcripts showed differential usage in one exon (Fig 6).

**Figure 5.**
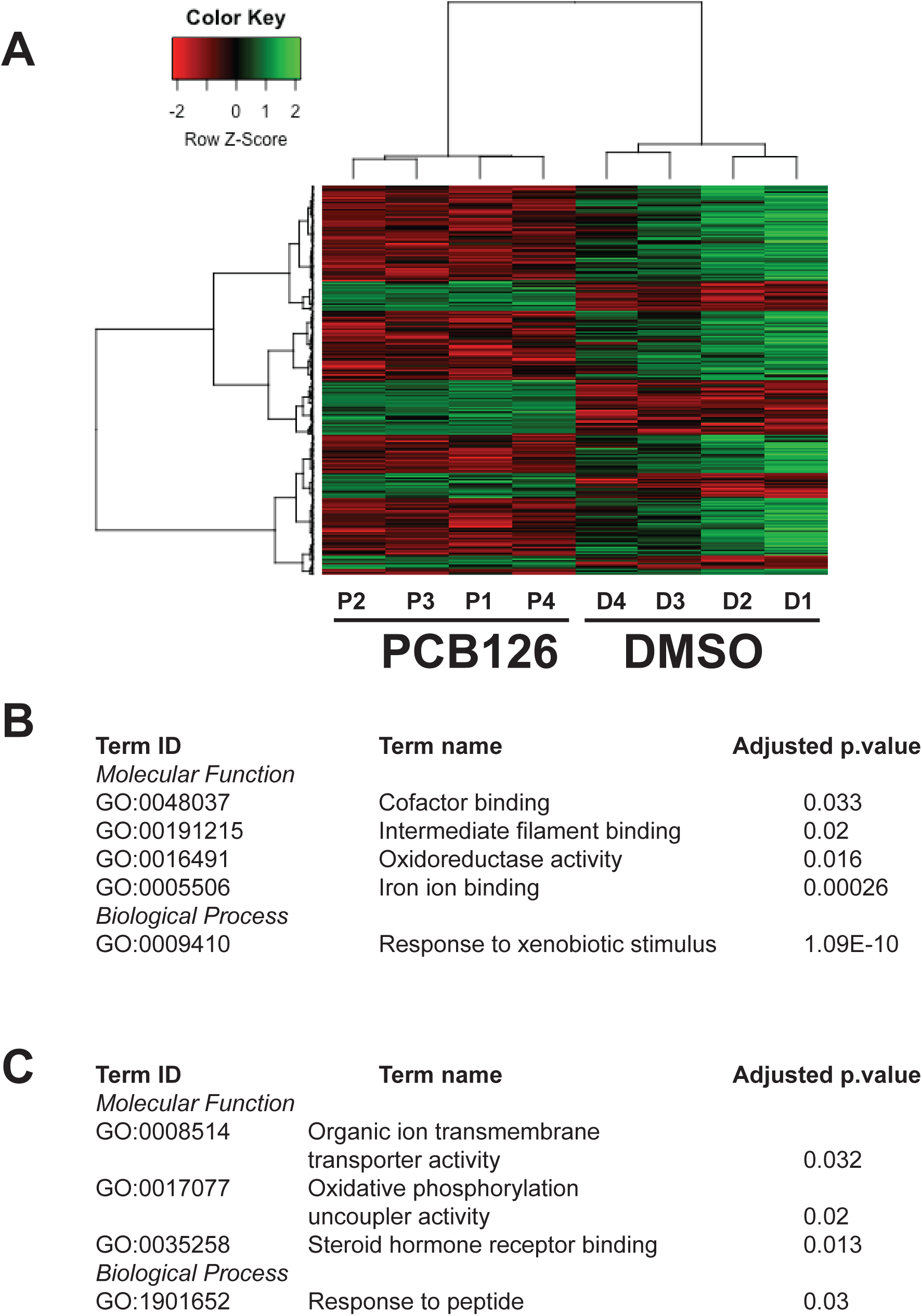
PCB126-induced differential gene expression. (A) Heatmap representation of the differentially expressed genes (DEGs). Differential gene expression analysis was performed on the input samples. (B and C) Gene Ontology (Biological Process and Molecular Function) terms of up and downregulated DEGs and their statistical significance (adjusted p.value <0.05).

**Figure 6.**
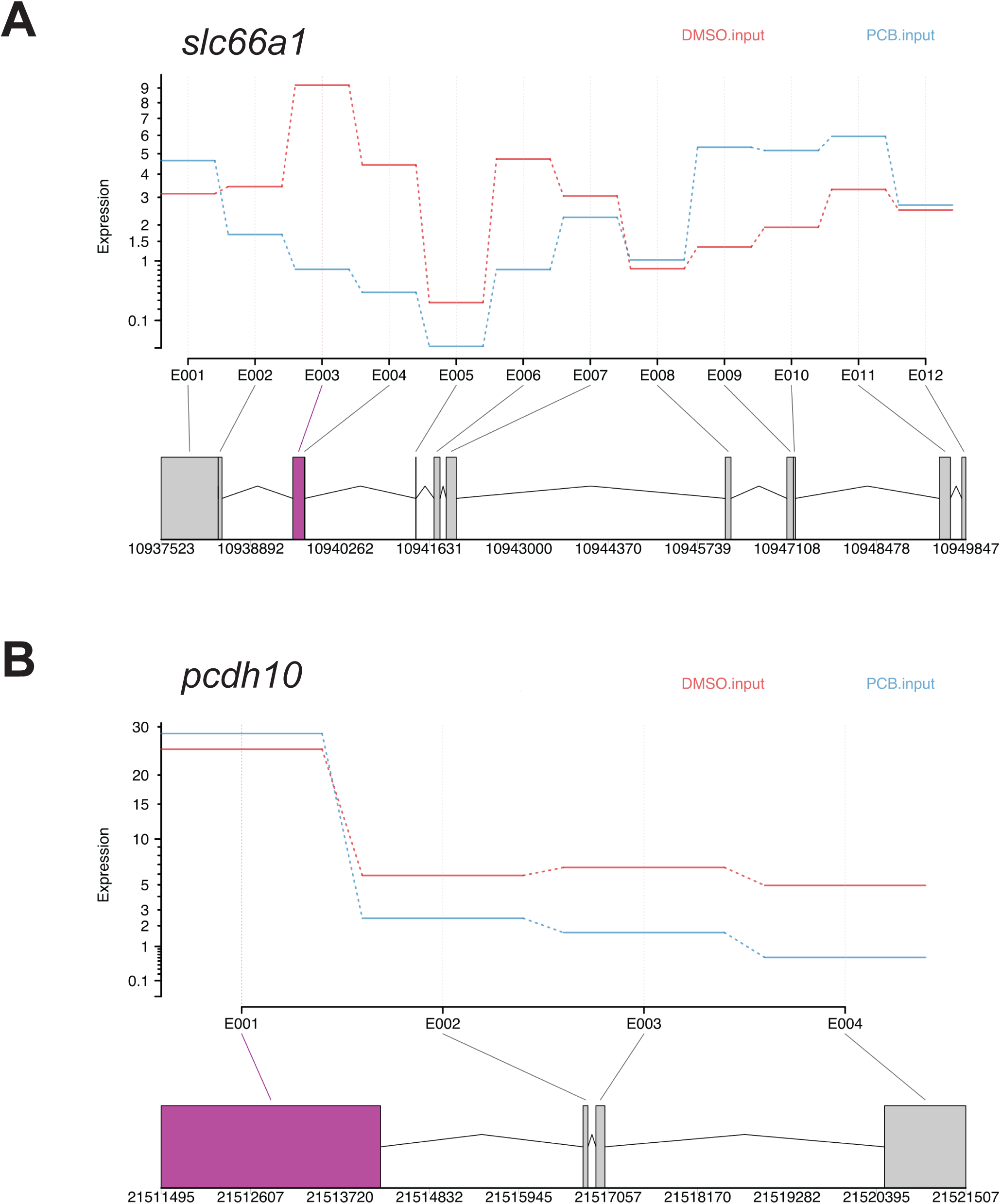
Effect of PCB126 exposure on differential exon usage. DEXSeq plots showing differential exon usage in *slc66a1* (A) and *pcdh10* (B) genes.

## Discussion

The present study demonstrated that acute PCB126 exposure alters m^6^A methylation of transcripts in zebrafish embryos. The functional significance of these alterations in developmental toxicity is not yet clear. However, based on the evidence that m6A RNA methylation plays a significant role during development (Heck and Wilusz 2019; Li et al. 2019; Vu et al. 2019), our results suggest that environmental chemicals can influence developmental processes by altering an important layer of post-transcriptional regulation of mRNAs and noncoding RNAs. Furthermore, m6A RNA methylation has been shown to play a key role in normal physiology and metabolism, stress response and in various types of cancer (Engel et al., 2018; Galardi et al. 2020; He et al. 2019), suggesting additional toxicity mechanisms for environmental chemicals.

Using MeRIP-seq, an unbiased transcriptome-wide approach, we demonstrated that acute exposure to PCB126 altered m6A methylation marks in coding and non-coding RNAs. These results provide the first evidence that m6A methylation marks can be altered by environmental chemical exposures. A recent study using an ELISA-based approach have shown that environmental chemicals such as bisphenol A, vinclozolin, sodium arsenite and particulate matter (PM2.5) affect m6A RNA methylation levels (Cayir et al., 2019). However, this approach does not show the location of the m6A alterations in the transcriptome. Using MeRIP-seq, we observed 15 transcripts to be differentially methylated in response to PCB126 exposure in zebrafish embryos. They include 11 transcripts with increased m6A methylation marks and 4 transcripts that lost methylation marks in response to PCB126 exposure. It is not clear how the gain or loss of m6A in response to PCB126 affects transcript fate, but recent studies have shown that m6A methylation influences RNA processing, export, translation or decay (Coots et al. 2017; Ke et al. 2017; Xiao et al. 2016). One of the best characterized mechanisms is the recruitment of proteins (readers) that recognize and bind to m6A modifications and how this affects RNA metabolism. One such group of m6A readers are the YTH domain containing family of proteins (e.g., YTHDF1-3, YTHDC1 and YTHDC3). These proteins preferentially bind to m6A-modified RNAs, recruit other effector proteins, and influence multiple aspects of RNA metabolism (Du et al. 2016; Wang et al. 2015). During maternal-zygotic transition, m6A RNA methylation through recruitment of YTHDF2 has been shown to induce decay of maternal transcripts in zebrafish (Zhao et al., 2017). A similar role for YTHDF2 has been observed in mammals (Ivanova et al. 2017). In addition, there is compelling evidence that several other proteins such as insulin-like growth factor 2 mRNA binding proteins (IGF2BPs) preferentially bind to m6A-modified RNAs. Further studies are needed to characterize the m6A-modified RNA-protein interactions in response to environmental chemical exposures.

Among the differentially methylated transcripts are products of three classical AHR target genes, *ahrra, tiparp* and *nfe2l2b. Ahrr and tiparp* are an integral part of AHR signaling and both have been shown to negatively regulate AHR activity by distinct mechanisms (MacPherson et al. 2014). In addition, recent studies demonstrated that *nrf2b*, a homolog of mammalian nuclear factor, erythroid 2-like 2 (NFE2L2) also act as a repressor of gene expression (Bahn et al. 2019; Liu et al. 2018; Thangasamy et al. 2011; Timme-Laragy et al. 2012). The expression of these transcripts was upregulated in response to PCB126 exposure. It remains to be determined how increased methylation affects expression of these transcripts. Previous studies have shown that m6A methylation affects mRNA splicing. For instance, knockout of m6A writers and/or eraser proteins affects alternative splicing (AS) patterns (Dominissini et al. 2012; Ping et al. 2014; Zhao et al. 2014; Zheng et al., 2013). Similarly, m6A reader proteins recruited to m6A-modified transcripts are also shown to cause AS events by recruiting splicing factors (Kasowitz et al., 2018). However, differential exon usage analysis did not reveal alternative splicing events among the differentially methylated transcripts. A recent study using exon arrays demonstrated that a potent AHR agonist, 2,3,7,8-tetrachlorodibenzo-p-dioxin (TCDD) induces alternative splicing in *ahrr* and *tiparp* transcripts in rodent livers (Villasenor-Altamirano et al. 2019). Whether these AS events are due to m6A RNA methylation remains to be determined. Similarly, further characterization of post-transcriptional regulation of AHR target genes in zebrafish embryos is necessary to determine the effect of m6A RNA methylation on alternative splicing during periods of cellular differentiation.

In addition to AHR signaling, PCB126 exposure hypermethylated a few other coding and noncoding transcripts. Interestingly some of these transcripts were previously shown to be associated with AHR signaling. One of these is *plekhh3*, whose expression is inducible by AHR agonists in zebrafish and in mammals (Fracchiolla et al. 2011; Frericks et al. 2006). Another related hypermethylated transcript was *plekhg4b*, with unknown function. Pleckstrin homology domains are present in a variety of modular proteins involved in cellular signaling, cytoskeletal organization, membrane trafficking and phospholipid processing (Scheffzek and Welti 2012). Even though the role of *plekhh3* in AHR-induced signaling has not been characterized, biochemical mechanisms that mediate cellular toxicity by causing cytoskeleton disruption have been demonstrated. We also observed hypermethylation of *rpgra* transcripts, which are primarily localized to microtubules. Studies in mammals have demonstrated complex alternative splicing patterns of the RPGR gene, with over 20 splice variants (Ferreira 2005; He et al. 2008). It is possible that AS is regulated by m6A RNA methylation. Mutations in *rpgra* are associated with atrophic macular degeneration and primary cilia dyskinesia (Bukowy-Bieryllo et al. 2013; Mawatari et al. 2019) and splicing mutations in AHR have been associated with the development of retinitis pigmentosa (Zhou et al. 2018). These results suggest that transcripts associated with AHR signaling are regulated by the m6A RNA modifications. Interestingly, the expression of *plekhh3* and *plekh4gb* is upregulated in response to PCB126 exposure. Further studies are needed to demonstrate the role of the epitranscriptome in xenobiotic and physiological functions of AHR.

We also observed four transcripts that lost methylation marks in response to PCB126 exposure, which are all related to the nervous system (Balamurugan and Sterneck 2013; Lee et al. 2003; Lewis et al. 2015). For instance, *vgf*, a highly conserved neuroendocrine factor expressed exclusively in neuronal and neuroendocrine cells and induced by various growth factors (nerve growth factor, brain-derived neurotrophic factor and glial-derived growth factor) was hypomethylated in response to PCB126 exposure. VGF mRNA levels are regulated by neuronal activity, including long-term potentiation, seizure, and injury (Lewis et al., 2015). Similarly, the *cebpd* gene encoding the C/EBPδ protein, a member of the C/EBP transcription factor family (Balamurugan and Sterneck 2013) that modulates many biological processes including cell differentiation, proliferation, and cell death, was hypomethylated. Concomitantly, *vgf* and *cebpd* expression was also downregulated. In addition, forkhead transcription factor Foxi1, an important player in embryogenesis particularly in the inner ear development was hypomethylated. These results are not surprising given the fact that dioxin-like PCBs have been shown to affect the nervous system (Aluru et al. 2017; Glazer et al. 2016; Orito et al. 2007; Piedrafita et al. 2008). Several recent studies have shown that m6A RNA modifications play an essential role in nervous system development (Livneh et al. 2020; Xu et al. 2020; Yu et al. 2020). Further studies are needed to determine the role of mRNA m6A methylation in dioxin-like PCB effects on brain development, learning and memory and other nervous system processes.

Our results led us to hypothesize that toxicant-induced alterations in m6A levels regulate the expression of transcripts by affecting mRNA stability. This is based on the observation that out of the 15 differentially methylated transcripts, 8 of them were found to be differentially expressed. Biochemical characterization studies have demonstrated that m6A is selectively recognized by binding proteins such as YTH domain family proteins and regulate mRNA stability (Wang et al. 2014). It remains to be determined if toxicant-induced changes in gene expression are under epitranscriptomic regulation.

## Conclusions

Here, we describe the first *in vivo* evidence demonstrating the effect of exposure to environmental chemicals on m6A RNA methylation landscape during vertebrate development. Our results indicate that regulation of m6A RNA methylation levels is one of the mechanisms by which PCB126 alters gene expression patterns. Further studies are necessary to uncover the role of AHR in the regulation and biological functions of m6A during zebrafish development. In addition, characterization of the role of m6A RNA methylation in toxicant-induced phenotypes will provide a better understanding of the range of pathologies and diseases associated with environmental chemical exposures (Yang 2020). Ever since the development of the MeRIP approach, substantial progress has been made in profiling m6A RNA modifications in a number of model systems. However, considerable limitations exist in quantitative determination of m6A patterns (McIntyre et al., 2020). Recent methodological developments have overcome some of these hurdles and hence paving the way for better understanding of this important layer of gene regulation (Linder et al., 2015; Zhang et al., 2019; Meyer et al., 2019).

## Supporting information

Supplemental spread sheet_DGE

Supplemental spread sheet_m6Apeaks

## Acknowledgements

This work was supported by the National Institute of Health Outstanding New Environmental Scientist Award to NA (NIH R01ES024915) and Woods Hole Center for Oceans and Human Health [National Institutes of Health (NIH) grant P01ES021923 and National Science Foundation Grant OCE-1314642 to M. E. Hahn, J. J. Stegeman, NA and SK]. The authors would like to thank the toxicology lab group members at WHOI for their suggestions with data interpretation.

