## Supplementary figures and images for "PCB126 exposure revealed alterations in m6A RNA modifications in transcripts associated with AHR activation"

### Supplemental Figure 1.

Supplementary Figure 1.

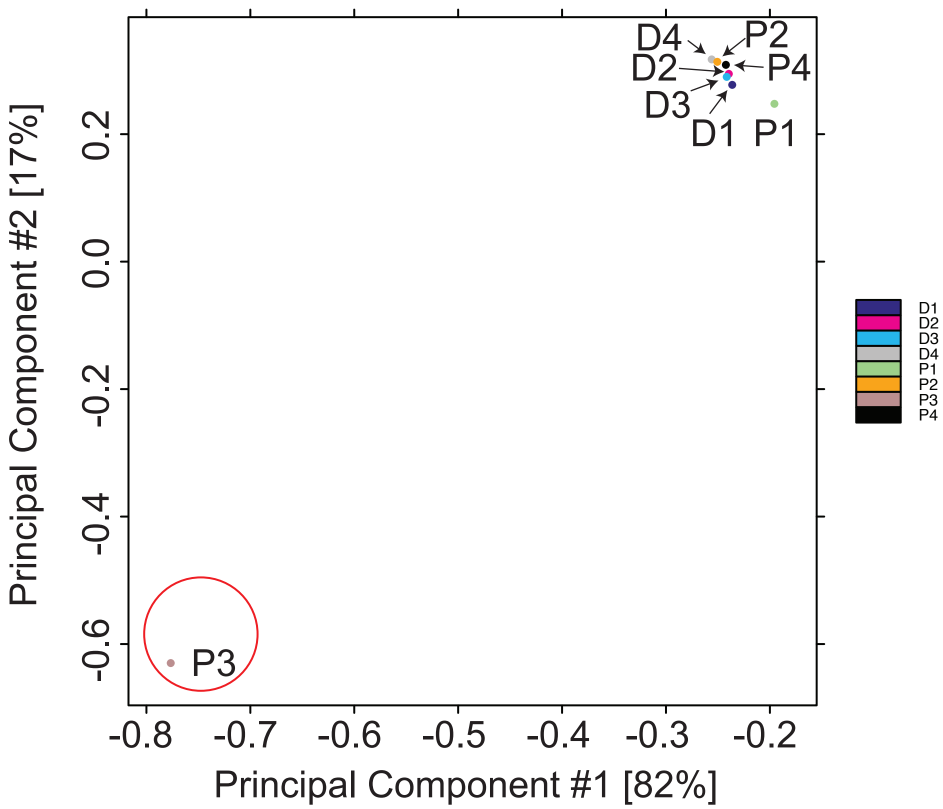
